# Porcine epidemic diarrhea virus Whole-Genome Diversity Reveals Divergence of Contemporary Populations in the United States

**DOI:** 10.64898/2026.10.08.757685

**Authors:** Joao P. H. da Silva, Albert Rovira, Fabio Vannucci, Juan Mena, Nakarin Pamornchainavakul, Melanie Prarat Koscielny, Jonah Perkins, Qirui Zhang, Mariana Kikuti, Xiaomei Yue, Igor A. Paploski, Montserrat Torremorell, Cesar Corzo, Kimberly VanderWaal

## Abstract

Porcine epidemic diarrhea virus (PEDV) was introduced into the United States (USA) in 2013 and has circulated ever since. PEDV causes significant losses in the swine industry, with the greatest impact observed during the first years following its introduction into the country. However, the evolutionary dynamics of the virus remain underexplored. Recently, analyses of the Spike protein subunit S1 suggested that the PEDV population currently circulating in the USA represents a fraction of its historical diversity and comprises only two clades restricted to specific geographic regions. These observations were based on a limited set of S1 sequences. Furthermore, our understanding of evolutionary dynamics at the genomic level is even more limited, as complete PEDV genomes have not been generated since 2017. Here, we characterized the evolutionary dynamics of PEDV populations in the USA using complete genomes and assessed the contribution of different evolutionary mechanisms to shaping population structure. We analyzed 338 complete genomes, of which 25 were obtained in this study, together with 623 Spike S1 sequences, of which 41 were obtained in this study. All sequences obtained in this study were assigned to the non-S-INDEL strain. No S-INDEL sequences were recovered in this study, although these viruses continue to circulate in the country. Our analyses revealed that most sequences sampled after 2023 formed a monophyletic clade divergent from sequences collected in previous years, suggesting an ongoing temporal population structuring. Population structure analyses further supported evidence of population differentiation over time. Recombination analyses indicated that sequences belonging to the contemporary clade shared a single recombination event, suggesting that this clade may have emerged through recombination. Time-scaled S1 phylogenies were consistent with previous findings, supporting the circulation of only two contemporary clades; however, one of these clades was detected for the first time outside its previously reported geographic range. Our findings document temporal genetic differentiation in US PEDV populations, consistent with continued viral evolution, and broaden the known geographic distribution of contemporary S1 clades. These results highlight the value of continued genomic surveillance to monitor changes in PEDV diversity and distribution and provide sequence data to inform vaccine research.

## Introduction

Over the last decade, porcine epidemic diarrhea virus (PEDV) eradication has been increasingly discussed in the United States. A key factor in the success of any disease eradication program is the identification and characterization of viral genetic diversity (1, 2). Understanding the processes that shape the genetic structure of viral populations across space and time provides insights into lineage emergence, persistence, turnover, extinction, dispersal, and adaptation (2–4). This information is fundamental for identifying risks and mitigating them through more effective surveillance, biocontainment, and vaccine development strategies.

PEDV, classified under the specie *Alphacoronavirus porci* within the family *Coronaviridae*, is a positive-sense single-stranded RNA virus with a genome of approximately 28 kb that causes acute enteric disease in pigs, resulting in significant economic losses to the global swine industry (5, 6). The virus was introduced into the United States in 2013 and has circulated in USA swine herds ever since (7, 8). The PEDV epidemic in the United States is typically divided into two periods: the epidemic period (April 2013 to August 2014) and the endemic period, which extends to the present day (9). Two distinct strains are known to circulate in USA swine populations: the S-INDEL (low pathogenic) and non-S-INDEL (high pathogenic) strains, which are characterized by the presence or absence of insertions and deletions in the S1 subunit region of the Spike protein, respectively (8, 10–12).

Most studies on PEDV population dynamics in the USA were conducted during the epidemic period, which coincided with more intensive sequencing efforts (7, 8, 13). Subsequently, the number of sequences generated per year decreased drastically, with a median of approximately 2.2 Spike subunit 1 (S1) protein sequences generated for every ten reported outbreaks (13). When complete genomes are considered, the limitation is even greater, as the most recent documented sequences are from 2017, resulting in a large gap in genomic information. Despite this sampling limitation, a recent study based on Spike (S1) sequences suggested that the composition of the viral population circulating in the United States has changed over time (13). Many of the clades circulating during the epidemic period were no longer detected after 2017, while only two clades descended from lineages present during that period continued to diversify and persist in the population. In addition, these two clades were found to be geographically structured, suggesting limited gene flow between the regions where contemporary viral populations circulate (13).

Although the findings of Herrera da Silva et al. (13) contributed to advancing our understanding of PEDV population dynamics, substantial sampling gaps remain, generating uncertainty regarding the extinction of some older clades as well as the current geographic structure of the viral population. In addition, most analyses in that study focused on the S1 subunit of the Spike protein due to the scarcity of recent complete genomes, preventing a comprehensive assessment of contemporary genetic diversity across the entire viral genome (13). To address these gaps, we collected samples from PEDV-infected pigs across the four major swine-producing regions of the United States. Our objectives were to confirm the geographic distribution of the two contemporary clades, evaluate whether the clades circulating during the epidemic period have indeed become extinct, and characterize the evolutionary trajectories of contemporary PEDV populations using complete genome sequences. These findings provide important information to guide biocontainment strategies, identify genomic regions that may represent potential targets for vaccine development, and contribute to ongoing PEDV control and eradication efforts in the United States.

## Methods

### Sampling

A total of 46 fecal swab samples were collected from PEDV-positive farms experiencing clinical outbreaks between 2017 and 2026. The target number of samples per region was defined to reflect that region’s proportion of the number of outbreaks reported to the Morrison Swine Health Monitoring Project (MSHMP; private data). Accordingly, 20 samples were collected from the Atlantic Seaboard, 15 from the Midwest, 6 from the Northeast, and 5 from the Southwest. For confidentiality reasons, the geographic distribution of the sequences is not explored further in this study. Of the 46 samples, 23 complete genomes were generated in this study. These include 6 genomes from the Atlantic Seaboard, 8 from the Midwest, 6 from the Northeast and three from the Southwest. Two additional complete genomes were provided by the Ohio Department of Agriculture Animal Disease Diagnostic Laboratory. Together, these 25 genomes represented samples collected between 2017-2018 and 2024-2026 (Table 1). No genomes from 2019-2023 were generated in this study. In addition, 35 Spike subunit 1 (S1) sequences were provided by collaborators, including 30 sequences from the Torremorell Laboratory (2017–2018, samples collected in the Midwest) and five sequences from the Ohio Department of Agriculture Animal Disease Diagnostic Laboratory (2017–2023, samples collected in the Northeast). We also incorporated 313 complete genome sequences and 565 S1 sequences from the USA previously analyzed by Herrera da Silva et al. (2025) (13) (Table 1).

**Table 1.** Temporal distribution of complete genome and Spike (S1) sequences included in this study. Values in parentheses indicate sequences generated in this study. Single asterisks (*) indicate that all Spike (S1) sequences generated in this study for that year were derived from complete genomes. Double asterisks (**) indicate that some, but not all, Spike (S1) sequences generated in this study for that year were extracted from complete genomes. Therefore, the complete genome and Spike (S1) datasets are not mutually exclusive. NA, not available.

| Year | Whole Genomes (n) | Spike (n) |
| --- | --- | --- |
| 2013 | 112 | 112 |
| 2014 | 153 | 296 |
| 2015 | 36 | 55 |
| 2016 | 7 | 22 |
| 2017 | 6 (1) | 33 (11**) |
| 2018 | (1) | 41 (31**) |
| 2019 | NA | 15 |
| 2020 | NA | 5 |
| 2021 | NA | 5 |
| 2022 | NA | 7 |
| 2023 | NA | 9 (1) |
| 2024 | (5) | 13 (5*) |
| 2025 | (17) | (17*) |
| 2026 | (1) | (1*) |
| Total | 338 (25) | 623 (60**) |

### RNA extraction

Fecal swab samples were collected from animal showing clinical. Samples were re-suspended by swirling in 1 mL of PBS (Thermo Fisher Scientific, Waltham, MA), followed by total RNA extraction using the MagMAX™ Viral RNA Isolation Kit (Life Technologies, Carlsbad, CA) on a KingFisher magnetic particle processor (Thermo Fisher Scientific, Waltham, MA), following the manufacturer’s recommendations with minor modifications. Briefly, RNA was eluted in 50 μL of elution buffer. Extracted nucleic acids were subsequently treated with RNase-free DNase I (Qiagen, Hilden, Germany) for 15 min at ambient temperature and then purified using Agencourt® RNAclean® XP Beads (Beckman Coulter Life Sciences, Indianapolis, IN) according to the manufacturer’s instructions.

### Library preparation and sequencing

Reverse transcription was performed using total RNA as the template. The cDNA was purified with Agencourt® AMPure® XP Beads (Beckman Coulter Life Sciences, Indianapolis, IN) per manufacturer’s recommended procedure and was re-suspended into 9 μl re-suspension buffer. The concentration of cDNA was determined by Qubit® 2.0 Fluorometer and Qubit® dsDNA HS Assay Kit (Thermo Fisher Scientific, Waltham, MA) and adjusted to 0.2 ng/μl for sequencing library preparation. Sequencing library was prepared using Nextera® XT library preparation kit (Illumina, San Diego, CA) with dual-indexing and sequencing was performed on an Illumina MiSeq® instrument (Illumina, San Diego, CA) according to user manual of “MiSeq Sequencing System Guide” for 2□×□150 base pair (bp) reads.

### Sequencies assembling

Initially, the raw reads were subjected to quality assessment and adapter screening using FastQC v0.12.1 (14). Reads were trimmed, and low-quality bases were removed using a Phred quality threshold of Q30 (15). Genome assembly was performed de novo using SPAdes genome assembler v4.2.0. To estimate coverage and sequencing depth, the reads were mapped against the assembled contigs using Bowtie2 v2.5.3 (16), and coverage statistics were calculated using Samtools v1.22 (17). The alignment of reads to the assembled contigs was visually inspected using Tablet genome viewer v1.21.02.08 (18).

### Multiple sequence alignments and phylogenetic analyses

Multiple sequence alignments were performed using the MAFFT algorithm v7.490 with default parameters for both complete genomes and Spike (S1) sequences (19). For the complete genome alignments, the 5′ and 3′ untranslated regions (UTRs) were trimmed, retaining only the coding regions for subsequent analyses. Initially, sequences were classified as S-INDEL or non-S-INDEL based on the presence or absence of specific deletions in the spike gene.

Phylogenetic trees were reconstructed for both complete genomes and Spike (S1) sequences using the maximum-likelihood approach implemented in RAxML-NG(20), with 1000 bootstrap replicates, under the best-fit nucleotide substitution model general time-reversible with a proportion of invariant sites and gamma-distributed rate variation among sites discretized into four categories (GTR+I+G4) determined using ModelTest-NG (21). Phylogenetic trees were visualized using FigTree v1.4.4 (http://tree.bio.ed.ac.uk/software/figtree).

Time-scaled phylogenetic trees were reconstructed using BEAST 1.10.4 (22) for the Spike (S1) after removing sequences showing evidence of recombination (see below). Due to the large number of recombination events detected throughout the complete genomes, we did not reconstruct time-scaled phylogenetic trees for the complete genome dataset, since the presence of these recombination events could distort parameters such as nucleotide substitution rates and time to the most recent common ancestor (TMRCA). Because no S-INDEL strains were identified in the newly generated dataset, no additional information could be obtained beyond that previously reported by Herrera Silva et al. (13). Therefore, only non-S-INDEL sequences were included in the time-scaled analyses.

To assess the evidence of temporal signal and determine the best-fitting molecular clock model, we estimated Bayes factors for competing models based on marginal likelihood estimation (23–25). To evaluate temporal signal, datasets were analyzed under heterochronous and isochronous sampling schemes. For molecular clock model comparison, we contrasted a strict molecular clock model and an uncorrelated relaxed lognormal molecular clock model. Time-scaled phylogenetic trees were reconstructed using the general time-reversible (GTR) substitution model with gamma-distributed rate heterogeneity across four categories under an exponential growth coalescent model.

Log marginal likelihood values were estimated using generalized stepping-stone sampling (GSS) (26). We used a Markov chain Monte Carlo (MCMC) chain length of 1 × 10□steps, with parameters logged every 1 × 10□steps. For the GSS analyses, we employed 100 path steps with a chain length of 1 × 10□iterations for each power posterior. Simulations were performed in two independent, and the convergence was assessed based on chain mixing and effective sample size (ESS) values (>200) using Tracer v1.7.3 (27). The runs were and combined using LogCombiner.

### Recombination analysis

The whole genomes were screened for evidence of recombination was screened using the RDP (28), GENECONV (29), BOOTSCAN (30), Maximum Chi Square (MAXCHI) (31), CHIMAERA (32), SisterScan (SiScan)(33), and 3SEQ (34) methods implemented in RDP5 v.5.84 (35). Statistical significance was assessed using a Bonferroni-corrected significance threshold of α = 0.05. Following the recommendations of the software authors, only recombination events supported by at least four independent methods were considered reliable (35).

For whole-genome sequences, phylogenetic incongruences potentially associated with recombination were further investigated using phylogenetic network approaches to capture signals of lateral transfer among genomic regions. Phylogenetic networks were inferred using the Neighbor-Net algorithm implemented in SplitsTree v.4.19.2 (36). Evidence of recombination was also assessed using the pairwise homoplasy index (PHI) test implemented in SplitsTree.

### Selection analyses

Coding regions of the PEDV genome were screened for evidence of positive and negative selection using three distinct methods: Single Likelihood Ancestor Counting (SLAC)(37), Mixed Effects Model of Evolution (MEME) (37), and Fast Unconstrained Bayesian Approximation (FUBAR) (38). All analyses were performed using the nucleotide substitution model that best fit each dataset as determined by the DataMonkey server and implemented through the DataMonkey webtool (39). Gene-wide nonsynonymous-to-synonymous substitution rate ratios (ω = dN/dS) were estimated for each coding region using the SLAC method. To account for the potential effects of recombination on selection inference, recombination breakpoints were identified using the Genetic Algorithm for Recombination Detection (GARD) (40), and the selection analyses were performed using the partition-specific phylogenetic trees inferred by GARD.

### Variability and Genetic structure of PEDV population in USA

For whole-genome sequences, the nucleotide diversity index, π (41), was estimated by computing the average number of nucleotide differences per site between pairs of sequences using a custom script written in R (available upon request) (42). Differences among mean π values were assessed by estimating 95% confidence intervals through non-parametric bootstrap tests with 10,000 replicates using the simpleboot (43) statistical package in R software (44). Nucleotide diversity was also estimated along the complete genome sequences using a sliding window approach with a window size of 200 nucleotide and a step size of 25 nucleotide. The number of segregating sites (S), number of haplotypes (H), and haplotype diversity (Hd), Watterson’s estimator of the population-scaled mutation rate (θW), and Tajima’s D were estimated using DnaSP v5.10 (45). Pairwise nucleotide identity was calculated using the Sequence Demarcation Tool (SDT) for whole-genome sequences, with comparisons performed within and between the clades defined in this study, including the contemporary and earlier non-S-INDEL groups (46).

We used a Mantel test to assess whether genetic distances between whole-genome sequences were correlated with temporal distances (47). Genetic distances were calculated as the number of nucleotide differences between each pair of sequences using the dist.dna function in the ape package in R (48), whereas temporal distances were calculated as the absolute differences between collection years. Pearson’s correlation coefficient was calculated between the two distance matrices, and its statistical significance was assessed using 99,999 permutations with the mantel function in the vegan package in R (49).

Population genetic structure was inferred using Discriminant Analysis of Principal Components (DAPC) implemented in the adegenet package in R (50, 51). DAPC was selected because it is a model-free, non-parametric approach that does not assume Hardy-Weinberg or linkage equilibrium among loci, unlike methods such as STRUCTURE (50, 51).The optimal number of clusters (K) was determined using the find.clusters function, which implements the k-means clustering algorithm across increasing K values. To determine the optimal number of clusters describing the complete genome dataset, k-means clustering was performed testing K values ranging from 2 to 10 while retaining all principal components (PCs). The optimal number of clusters was selected based on the inflection point of the Bayesian Information Criterion (BIC) curve, corresponding to the point at which further decreases in BIC became marginal (51). In addition, the genetic differentiation among the clusters inferred by DAPC was assessed using an analysis of molecular variance (AMOVA) implemented in the pegas R package (52) and by estimating the nucleotide differentiation coefficient (NST) in DnaSP v6 (45). To identify nucleotide sites contributing to the temporal genetic structure of PEDV populations in the United States, we inspected the variable loadings associated with each DAPC discriminant function using the adegenet R package (51, 53). The contribution of each nucleotide site was plotted for each discriminant function, and sites with the highest loading values were retained as candidate contributors to population differentiation. Specifically, informative sites were selected using the 99.5th percentile of the loading distribution for each discriminant function rather than by applying a fixed loading cutoff. For the retained sites, allele frequencies were then estimated and plotted across the corresponding DAPC groups to visualize how nucleotide states varied among groups.

## Results

### PEDV genomes sequence assembly and phylogenetic analysis

A total of 338 complete genomes and 623 Spike subunit 1 (S1) sequences originating from USA were analyzed. Of these 25 were complete genomes and 41 spike (S1) were obtained in this study. For the genomes generated in this study, coding regions showed 100% of coverage, with a minimum mean read depth of 100×. The sequences analyzed here cover a 13-year period (2013–2026). However, the complete-genomes dataset contains a gap in sequence availability between 2019-2023 (Table 1). In terms of geographic coverage, the dataset represents 16 USA states, spanning the four major swine-producing regions of the USA (Southwest, Midwest, Northeast, and Atlantic Seaboard). To preserve the anonymity of study participants who generously provided samples for this study, the geographic origin of individual samples is not explicitly reported.

A maximum-likelihood phylogenetic tree reconstructed using complete PEDV genomes revealed a clear phylogenetic separation between S-INDEL and non-S-INDEL strains (Figure 1). Within the non-S-INDEL strain, sequences exhibited a marked temporal structure. Most genomes collected before 2018 were placed in basal positions in the tree close to the root (hereafter referred to as early non-S-INDEL strains), whereas the majority of genomes sampled from 2024 onward clustered within a distinct, divergent monophyletic clade (recent non-S-INDEL strains). However, two genomes collected in 2025 grouped with the early non-S-INDEL strain rather than with the contemporary clade, suggesting that viruses phylogenetically related to the early non-S-INDEL strain are still detected in the current population.

**Figure 1.**
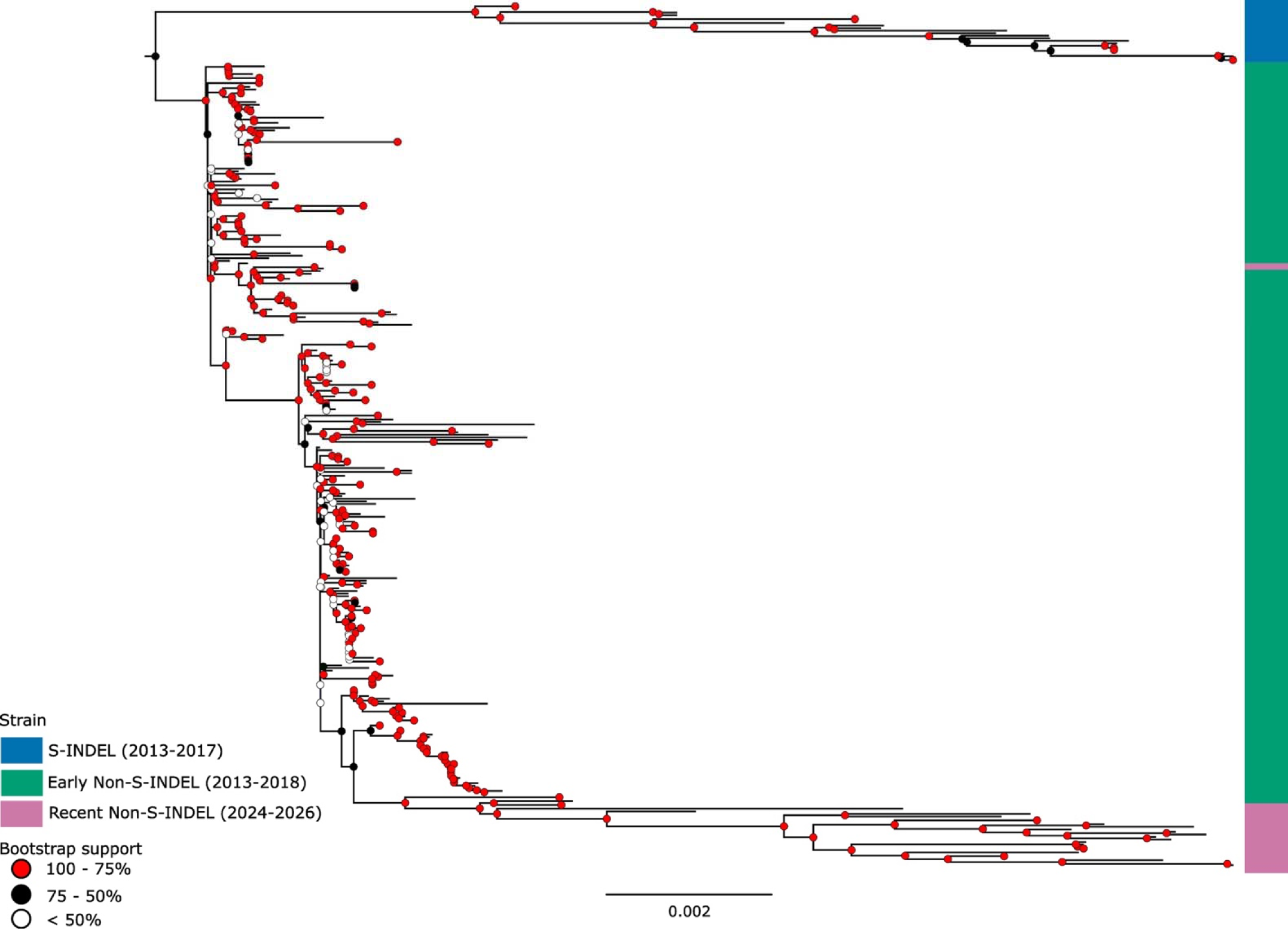
Midpoint-rooted maximum likelihood phylogenetic tree based on complete genome sequences. The phylogeny was reconstructed using RAxML-NG under the GTR+I+G4 nucleotide substitution model selected by ModelTest-NG. The colored side bars indicate strain classification, with S-INDEL strains shown in blue (colored side bar), early non-S-INDEL strains (2013–2017) shown in green, and recent non-S-INDEL strains (2018–2026) shown in pink. Nodes with bootstrap support values between 75% and 100% are shown in red values between 50% and 75% in black, and values below 50% in white. The scale bar represents the number of nucleotide substitutions per site.

In addition, we reconstructed a time-scaled phylogeny using recombination-free Spike protein subunit 1 (S1) sequences in BEAST. Because no S-INDEL sequences were detected in the present study, the time-scaled phylogeny was reconstructed only for the non-S-INDEL strain, as including S-INDEL sequences would not provide additional information beyond that previously described by Herrera da Silva (13). Our time-scaled phylogeny recovered the same overall population structure reported by Herrera da Silva (13), reinforcing previous evidence that most clades circulating during the epidemic phase are no longer detected, whereas only two major non-S-INDEL clades remain in circulation in the United States (Figure S1). Among the 60 S1 sequences generated in this study, 58 clustered within Clade 1, whereas only two grouped within Clade 2. These two Clade 2 viruses were the same two complete genomes that were closely related to the early non-S-INDEL strain described above (Figure S1).

Contrary to the previous observations that Clade 1 viruses were restricted to the Midwestern and Eastern United States and Clade 2 viruses to the Western United States (13), we detected a Clade 1 sequence sampled in the Western United States, suggesting that some level of PEDV dispersal occurs among these major geographic regions. Nevertheless, to date, Clade 2 viruses appear to remain restricted to the Western United States.

### Recombination analyses

Complete PEDV genomes were screened to evaluate evidence of intraspecific recombination. All 21 sequences belonging to the monophyletic clade that includes the recent Non-S-INDEL strain had a shared recombination event, event 13 (Figure 2C), covering the region from position 17,361 to 21,170, for which the major parental sequence was inferred to be unknown, whereas the minor parental sequence belonged to an early non-S-INDEL strain. This suggests that this clade likely emerged through recombination between an unknown major parental sequence and a minor parental sequence belonging to the non-S-INDEL strain derived from the earlier USA viruses (Table S1). In addition, six sequences within this contemporary clade also carried recombination event 9, spanning positions 4,545 to 16,815, which had previously been reported by Herrera da Silva et al. (13).

**Figure 2.**
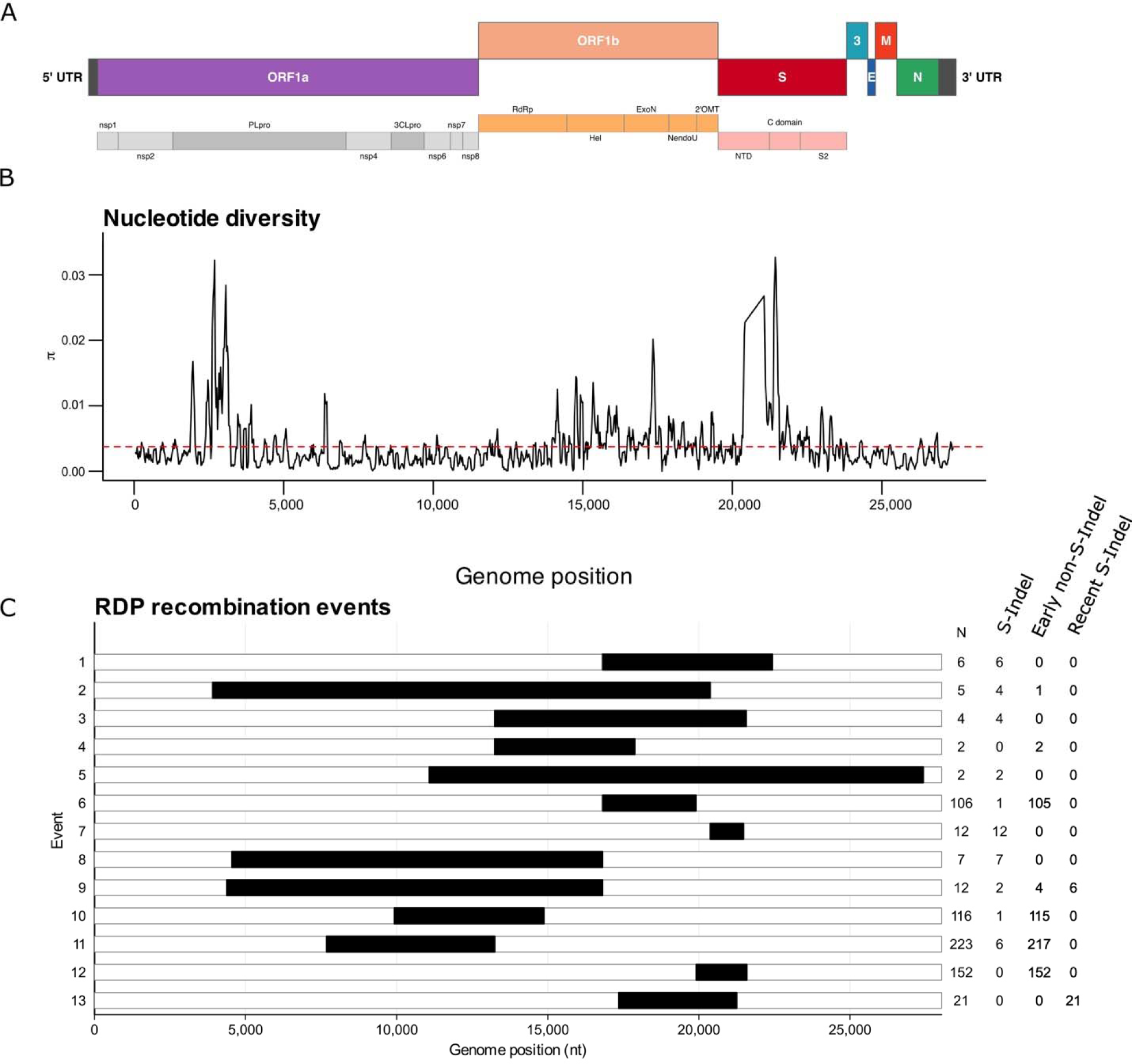
Genome-wide patterns of nucleotide diversity and recombination in PEDV. **A)** Genomic organization of PEDV, including the main ORFs and annotated protein domains. **B)** Nucleotide diversity (π) estimated across the genome using a sliding-window approach. The red dashed line represents the mean genome-wide nucleotide diversity. **C)** Recombination breakpoints inferred by RDP. Black segments indicate the genomic region involved in each recombination event. Numbers on the left indicate the recombination event ID, whereas numbers on the right indicate the number of sequences containing each respective event. On the right, N indicates the total number of sequences containing each event, whereas indicate the number of sequences assigned to the S-Indel, early non-S-Indel, and recent S-Indel groups, respectively.

The remaining 12 recombination events recovered in our analyses corresponded to those previously reported by Herrera da Silva et al. (13), as expected given the substantial overlap between the sequence datasets used in both studies (Figure 2).

Finally, we reconstructed a phylogenetic network to assess potential genealogical conflicts. While Herrera da Silva et al. (13) recovered four main groups, our phylogenetic network recovered five groups (Figure 3). One group corresponded to the S-INDEL strain, while the Non-S-INDEL strain was subdivided into four additional groups. Tree of these Non-S-INDEL groups were compact, with relatively short branch lengths, corresponding to isolates from the early Non-S-INDEL strain. Sequences belonging to the recent Non-S-INDEL strain formed a separate group, characterized by markedly longer branch lengths. However, two recent sequences did not follow this pattern and instead grouped with the early Non-S-INDEL isolates (indicated by the pink star in Figure 3), consistent with the exceptions noted in the whole-genome phylogeny.

**Figure 3.**
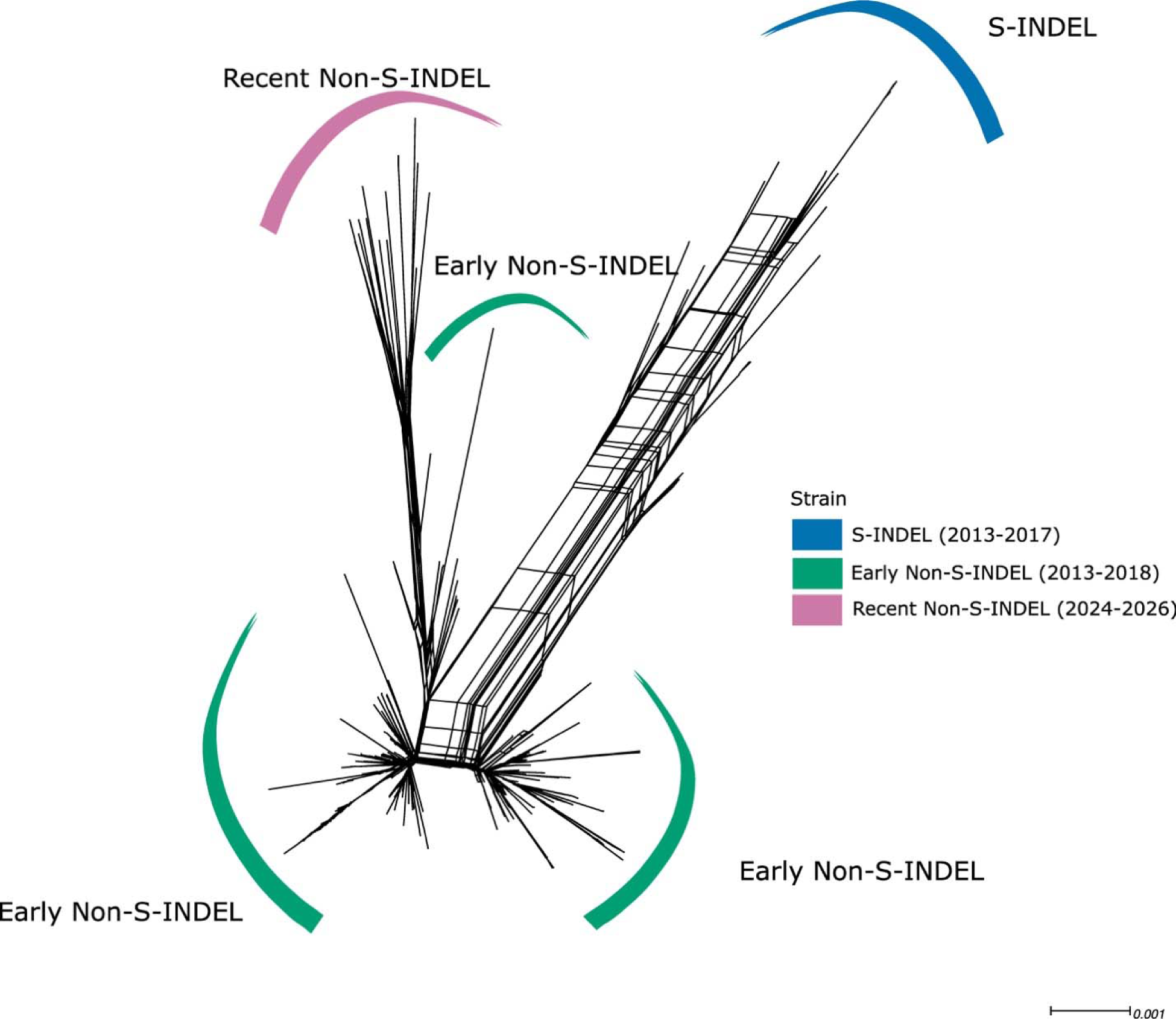
Phylogenetic network of PEDV whole-genome sequences. The network shows the genetic relationships among PEDV sequences based on the whole-genome alignment. Branch colors indicate the strain: S-INDEL, Early Non-S-INDEL, and Recent Non-S-INDEL. The reticulated structure indicates conflicting phylogenetic signals among sequences, consistent with the presence of recombination and/or shared polymorphisms across lineages. The scale bar represents nucleotide substitutions per site.

The reticulated patterns observed in the phylogenetic network provide evidence of phylogenetic conflicts potentially generated by recombination, with the S-INDEL branch exhibiting a denser reticulation pattern relative to Non-S-INDEL. The PHI test also suggested strong evidence of genealogical conflict among sequences (p > 0.0001). These findings are consistent with the recombination events detected by RDP: at least 10 distinct events were identified for the S-INDEL strain, compared with 7 for early Non-S-INDEL, of which 4 are shared by a massive fraction of the dataset (events 6, 10, 11, and 12), indicating their broad distribution. Interestingly, the recent Non-S-INDEL sequences that grouped within the divergent branch all shared the same recombination event, event 13, evidencing the impact of recombination on the diversification of this particular group.

### Genetic distance

Pairwise nucleotide identity was calculated using the whole genome sequences. The median pairwise identity among PEDV sequences was 99.58% (IQR 99.42–99.83%) (Figure 4). The median pairwise identity within the S-INDEL and non-S-INDEL strains was 99.51% (IQR 99.39–99.67%) and 99.77% (IQR 99.66–99.85%), respectively, whereas the median pairwise identity between these two strains was 98.93% (IQR 98.79–99.10%). Finally, we also estimated the genetic distances within the contemporary clade, which showed a median pairwise identity of 99.42% (IQR 99.34–99.53%), with a median pairwise identity of 99.15% (IQR 99.07–99.21%) relative to the remaining non-S-INDEL sequences collected during the earlier period and 98.61% (IQR 98.53–98.69%) relative to the S-INDEL strains.

**Figure 4.**
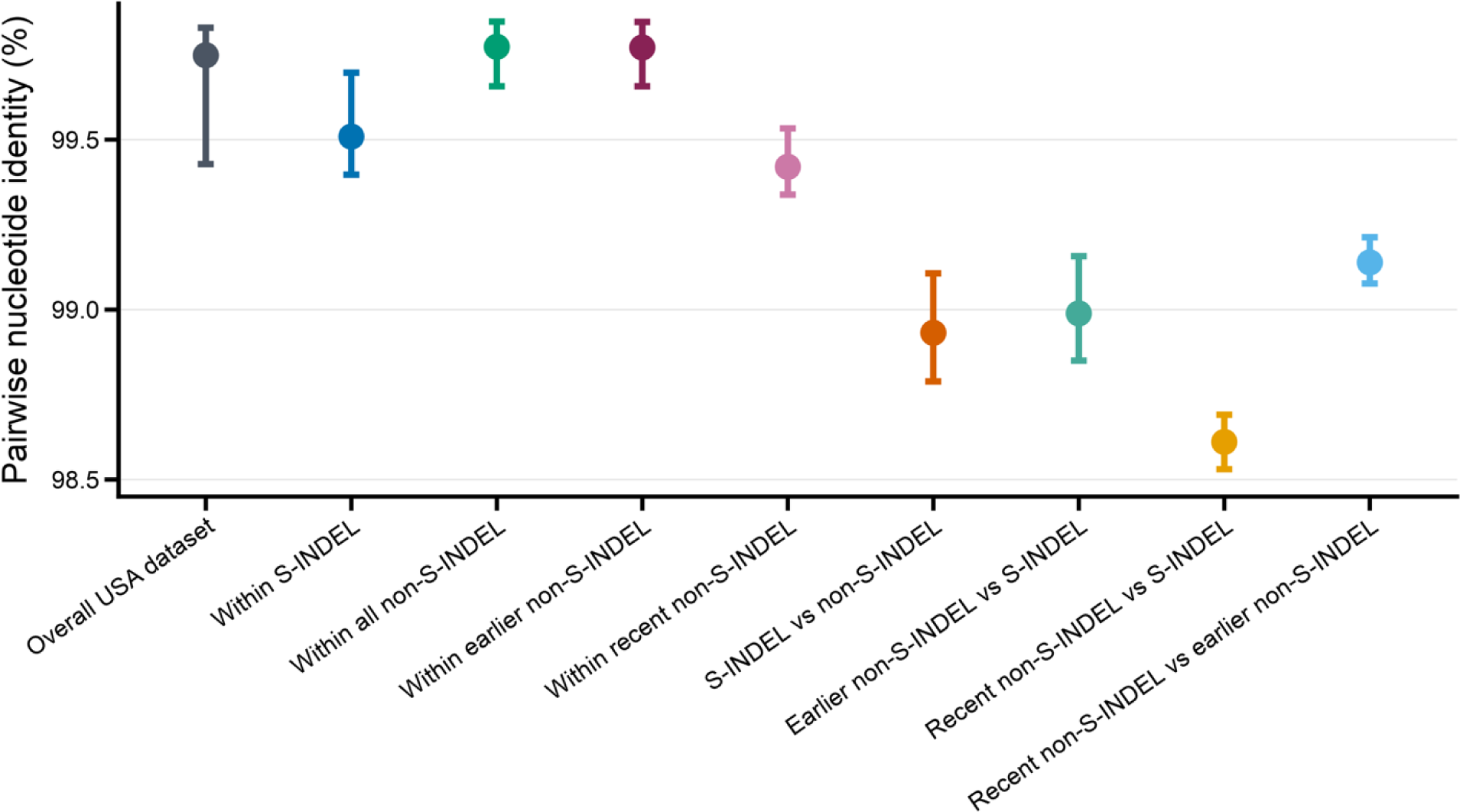
Pairwise nucleotide identity within and between PEDV groups. Pairwise nucleotide identity was estimated for the overall USA dataset and for within-and between-group comparisons among S-INDEL, early non-S-INDEL, and recent non-S-INDEL strains. Points represent median pairwise nucleotide identity, and error bars indicate the interquartile range (IQR). Lower values indicate greater genetic distance between groups.

### Population genetic structure of PEDV

Phylogenetic analyses together with the phylogenetic network, based on whole genome sequences, suggested patterns of population subdivision (Figure 1 and 3). In particular, the Non-S-INDEL strain displayed a subclade in which only isolates sampled after 2024 are grouped. To evaluate whether the genetic divergence between pairs of sequences was associated with sampling time, we performed a Mantel test, comparing the paired genetic distance and temporal distance matrices. The results of the Mantel test indicated a moderate, positive, and significant correlation between the matrices (r = 0.424; p = 1.0 × 10□□, 9,999 permutations), indicating that pairs of sequences sampled in more distant years tend to have greater genetic distance. These results suggest an ongoing process of temporal structuring for the PEDV sequences analyzed here (Figure 5).

**Figure 5.**
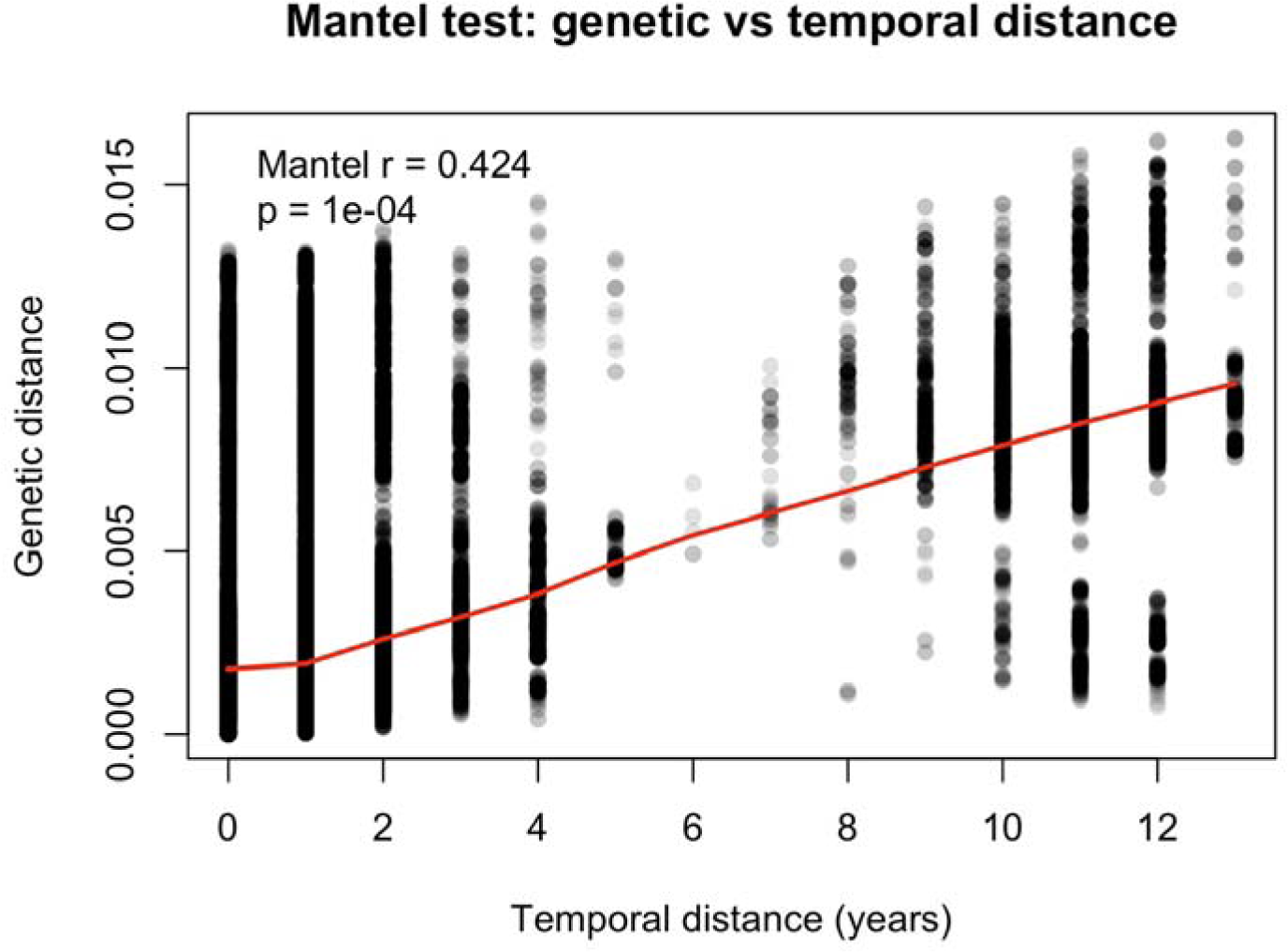
Association between genetic and temporal distances among PEDV sequences. Scatterplot showing pairwise genetic distance as a function of pairwise temporal distance between sequences. Each point represents a pairwise comparison between two sequences. The red line represents a smoothed trend fitted to the data, illustrating the overall increase in genetic distance with increasing temporal distance.

To access the population genetic structure of PEDV, we applied discriminant analysis of principal components (DAPC) to complete genomes using two complementary approaches. First, we performed an unsupervised DAPC to identify genetic clusters in multivariate space without assigning prior temporal categories to the sequences (Figure 6B). Next, we performed a second DAPC by grouping the sequences according to epidemiological period: initial epidemic period (2013–2014), post-epidemic period (2015–2018), and current period (2024–2026), with the objective of evaluating whether the observed genetic structure also reflected temporal patterns (Figure 6D).

**Figure 6.**
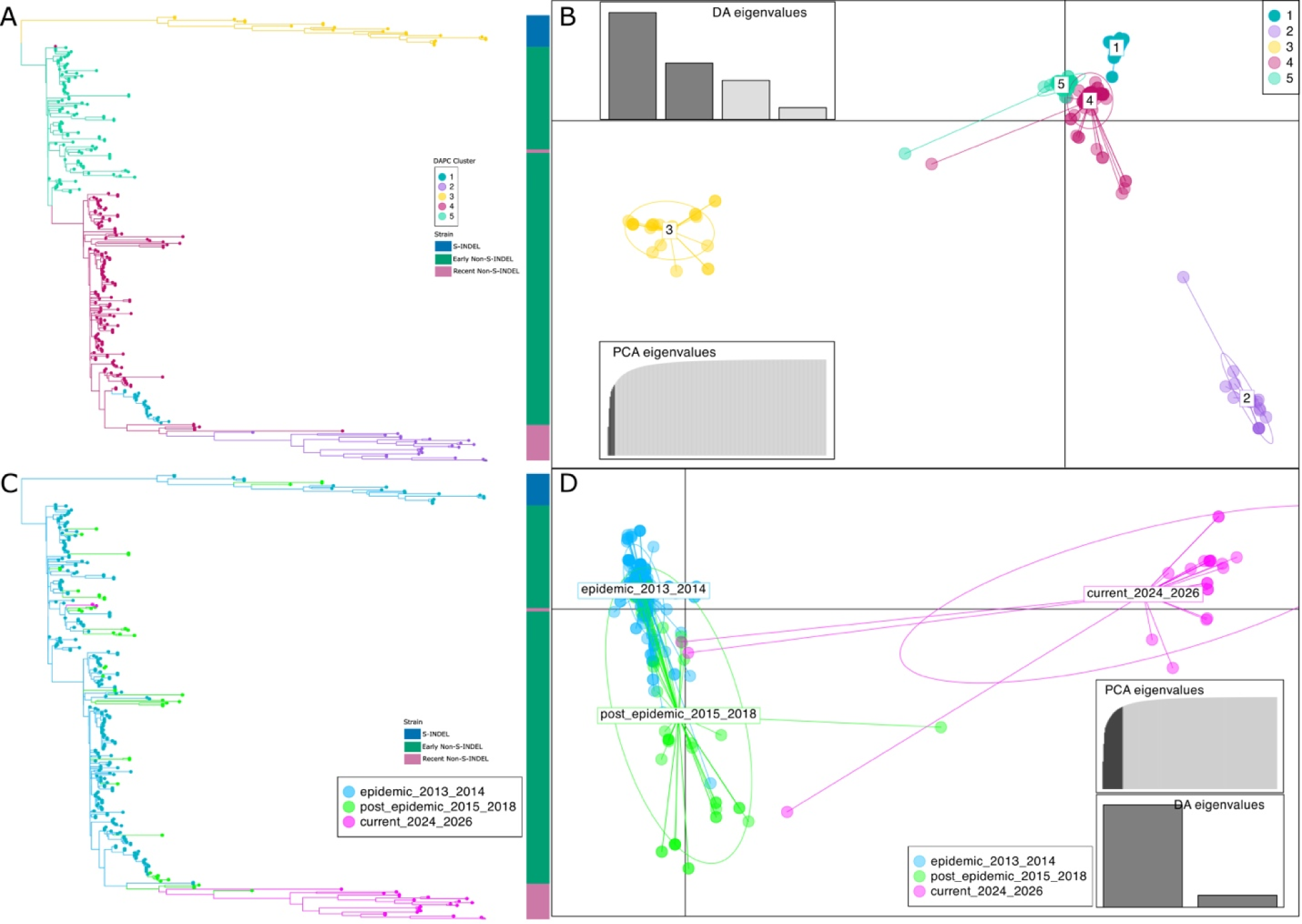
Genetic structure of PEDV populations. A. Midpoint-rooted maximum-likelihood phylogenetic tree, with colored circles at the tips representing the five genetic clusters inferred by DAPC. The lateral annotation bar indicates the strain classification corresponding to each clade. B. DAPC scatterplot based on the inferred genetic clusters inferred by DAPC; each point represents an individual sequence, and colors indicate the assigned DAPC cluster. The inset barplots show the eigenvalues retained by the discriminant analysis (DA eigenvalues) and by the principal component analysis (PCA eigenvalues). C. Midpoint-rooted maximum-likelihood phylogenetic tree with tip circles colored according to epidemiological period. The lateral annotation bar indicates the strain classification corresponding to each clade. D. DAPC scatterplot after stratifying sequences by epidemiological period; each point represents an individual sequence, and colors indicate the corresponding epidemiological period.

The unsupervised DAPC identified five major genetic clusters (Figure 6), which align well with the phylogenetic network. One of these clusters corresponded to the S-INDEL strain (Cluster 3), whereas the Non-S-INDEL sequences were subdivided into four distinct groups (Cluster 1, 2, 4 and 5) (Figure 6 A-B). Three of these groups were composed predominantly of sequences collected between 2013 and 2018 (Cluster 1, 4 and 5), although two sequences sampled in 2025 grouped within cluster 5. Interestingly, cluster 2 was composed exclusively of sequences sampled between 2024 and 2026 (Figure 6 A). The first two discriminant functions clearly separated clusters 2 and 3 from the others (Figure S2), whereas clusters 1, 4, and 5 were more closely positioned in the discriminant space, showing partial overlap but remaining distinguishable from one another (Figure S2).

To quantify the genetic structure observed in the DAPC, we performed an analysis of molecular variance (AMOVA). The AMOVA indicated strong differentiation among the five clusters inferred by the DAPC (ΦST = 0.839; p < 0.0001; 9,999 permutations), suggesting that a substantial proportion of the molecular variation is associated with differences among groups. To evaluate the magnitude of genetic differentiation between pairs of clusters, we estimated the nucleotide differentiation coefficient (NST). Pairwise NST values above 0.25 are indicative of genetic differentiation between groups. Pairwise NST values ranged from 0.119 to 0.653 (Table 2). Clusters 2 and 3 showed high differentiation relative to the others, whereas clusters 1 and 4 showed the lowest differentiation between each other. The 1–5 and 4–5 pairs showed intermediate differentiation (Table 2).

**Table 2.**
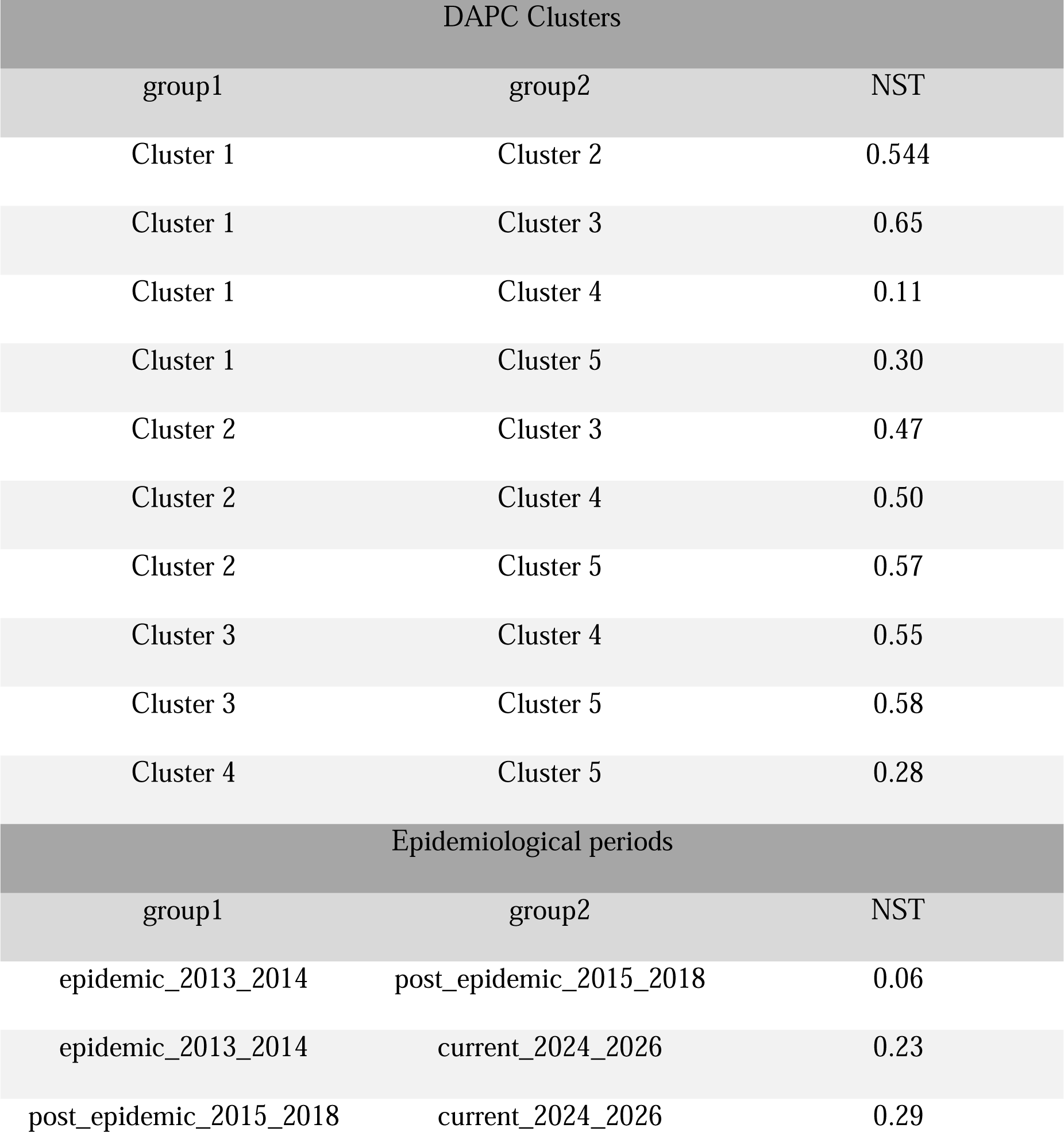
Pairwise nucleotide differentiation coefficient (NST) among DAPC-inferred clusters and epidemiological periods. NST values were estimated for all pairwise comparisons among genetic clusters inferred by DAPC and among epidemiological periods. Values above 0.25 indicate a higher degree of genetic differentiation between groups.

The DAPC stratified by epidemiological period revealed clear separation between current sequences (2024–2026) and those sampled before 2018, driven primarily by the first discriminant axis (Figure 6C–D; Figure S4), whereas substantial overlap remained between the initial epidemic and post-epidemic periods. A few contemporary sequences were positioned closer to the earlier populations, but most formed a distinct cluster. This pattern was corroborated by AMOVA (ΦST = 0.459; p < 0.0001; 9,999 permutations) of moderate magnitude compared to the unsupervised DAPC clusters and by NST values, which indicated moderate differentiation between the current period and earlier periods but very low differentiation between the epidemic and post-epidemic periods (Table 2).

### PEDV genetic variability

The genetic diversity of the complete PEDV genome dataset was evaluated. The 330 PEDV genomes analyzed here comprised 262 distinct haplotypes, corresponding to a haplotype diversity of 0.998 (Table 3). The nucleotide diversity (π) was substantially lower than Theta Watterson’s population-scaled mutation rate estimate (θW). Consequently, Tajima’s D was negative and significant, indicating an excess of low-frequency variants within the population. This pattern is compatible with recent population expansion following a bottleneck. However, because the complete dataset comprises genetically and temporally structured viral populations, this pattern may also be partially influenced by population structure (54, 55).

**Table 3.** Genetic diversity estimates for the complete PEDV dataset, DAPC-inferred clusters, and epidemiological periods. n, number of sequences; S, number of segregating sites; H, number of haplotypes; Hd, haplotype diversity; π, nucleotide diversity per site; θ, Watterson’s estimate of the population mutation rate per site based on the number of segregating sites; Tajima, Tajima’s D statistic. Asterisks indicate significant Tajima’s D values (p < 0.05).

| Group | n | S | H | Hd | pi | theta | tajima |
| --- | --- | --- | --- | --- | --- | --- | --- |
| Full dataset |  |  |  |  |  |  |  |
| Full dataset | 330 | 3024 | 262 | 0.998 | 0.0037 | 0.0172 | -2.44* |
| DAPC Clusters |  |  |  |  |  |  |  |
| 1 | 25 | 59 | 20 | 0.983 | 0.0003 | 0.00056 | -1.57 |
| 2 | 22 | 670 | 20 | 0.991 | 0.005 | 0.0067 | -0.89 |
| 3 | 22 | 435 | 19 | 0.982 | 0.004 | 0.0043 | 0.0019 |
| 4 | 153 | 1624 | 119 | 0.996 | 0.001 | 0.010 | -2.86* |
| 5 | 108 | 1350 | 80 | 0.994 | 0.001 | 0.0093 | -2.87* |
| Epidemiological period |  |  |  |  |  |  |  |
| Epidemic<br>2013-2014 | 258 | 1925 | 197 | 0.997 | 0.0027 | 0.0114 | -2.39* |
| Post<br>epidemic<br>2015-2018 | 48 | 1570 | 42 | 0.994 | 0.0042 | 0.0128 | -2.47* |
| Current<br>2024-2026 | 24 | 868 | 22 | 0.992 | 0.0063 | 0.0084 | -1.04 |

To account for population structure, we estimated the same diversity statistics for each of the DAPC-inferred genetic clusters. Clusters 2 (recent Non-S-INDEL) and 3 (S-INDEL) showed the highest nucleotide diversity (π = 0.005 and 0.004, respectively) compared with the other clusters (Table 3). However, Tajima’s D was not significant for either cluster, indicating no significant deviation from neutral model (Table 3). The Cluster 1 exhibited the lowest nucleotide diversity among all groups and showed a negative, although non-significant, Tajima’s D. Clusters 4 and 5 (both early Non-S-INDEL) displayed intermediate nucleotide diversity, with π values lower than θW and significantly negative Tajima’s D values, indicating an enrichment of low-frequency variants within these populations. The diversity patterns observed among the DAPC clusters were also consistent with branch lengths in the phylogenetic tree, with some clusters being composed of highly similar genomes, whereas others exhibited substantially greater genetic divergence.

To gain further insight into the demographic processes underlying the temporal population structure of PEDV in the United States, we next evaluated genetic diversity across the epidemic, post-epidemic, and contemporary epidemiological periods. The epidemic period (2013–2014) exhibited the lowest nucleotide diversity, with π substantially lower than θW and a significantly negative Tajima’s D. This pattern is consistent with the known epidemiological history of PEDV in the United States, in which the virus underwent a founder event upon its introduction into the country, followed by rapid population expansion (10, 11, 13). As expected under this demographic scenario, rapid expansion generates an excess of low-frequency variants, resulting in negative Tajima’s D values (54–57). During the post-epidemic period (2015–2018), nucleotide diversity increased approximately 1.5-fold relative to the epidemic period (Table 3).

Nevertheless, π remained lower than θW, and Tajima’s D remained negative, indicating that the excess of low-frequency variants persisted after the initial epidemic phase (Table 3). Interestingly, the contemporary period (2024–2026) exhibited the highest nucleotide diversity, approximately 2.3-fold higher than during the epidemic period and 1.5-fold higher than during the post-epidemic period (Table 3). In addition, π was more similar to θW, and Tajima’s D was not significant (Table 3).

Finally, we quantified nucleotide diversity across the complete PEDV genome (Figure 2B). Our results further reinforced our previous findings that the most variable genomic regions are located within ORF1a, particularly in the nsp2 and papain-like protease (PL-Pro) regions, as well as in the S1 portion of the Spike gene (13).

Taken together, these results indicate that the genetic diversity of PEDV has increased over time following its introduction into the United States. Despite this increase in diversity, viruses phylogenetically related to the early non-S-INDEL strain continue to be detected in the contemporary population.

### Selection analysis

To investigate how natural selection acts across the different coding regions of the PEDV genome, we performed selection analyses separately for each coding region. We estimated the ratio of nonsynonymous to synonymous substitutions (ω = dN/dS) for ORF1a, ORF1b, ORF3 and the ORFs encoding the spike (S), envelope (E), membrane (M), and nucleocapsid (N) proteins. All PEDV genes exhibited ω values lower than 1 (Table 4; Table S2), indicating evidence of purifying selection across all coding regions. However, the magnitude of purifying selection differed among genes. Spike gene and Envelope gene exhibited ω values closest to 1 (0.877 and 0.750, respectively), indicating more relaxed purifying selection, whereas Nucleocapsid showed an intermediate degree of purifying selection compared with the other genes analyzed. In contrast, ORF1a, ORF1b, ORF3, and Membrane gene appeared to be under stronger purifying selection. Although all genes were predominantly under purifying selection, most contained at least one codon inferred to be under diversifying selection, with the exception of Membrane and Envelope (Table 4; Table S2). The codons inferred to be under diversifying selection within the structural proteins did not differ from those previously reported by Herrera da Silva *et al.*(13), with the most positive selected codons being found in the Spike (Table 4; Table S2). However, in that study, codon selection analysis was performed only for structural proteins of the genome. Here, we detected codons under diversifying selection in ORF1a. Five codons were identified by MEME, indicating diversifying selection, whereas sixteen codons were identified by FUBAR, indicating diversifying selection. Only codons 477, 1011, and 1431 were detected by both methods, all of which are located within the papain-like protease region (Table 4; Table S2). ORF1b presented only one codon under diversifying selection detected by MEME and other two codons under diversifying selection detected by FUBAR; however, none were identified by both methods. Finally, ORF3 contained a single codon under pervasive diversifying selection detected exclusively by FUBAR.

**Table 4.** Nonsynonymous to synonymous substitution ratios (dN/dS) and number of codons inferred to be under diversifying or purifying selection in each PEDV gene. Selection analyses were performed separately for each coding region using SLAC, MEME, and FUBAR. The dN/dS ratio represents the overall nonsynonymous to synonymous substitution rate estimated for each gene. Values below 1 indicate predominant purifying selection. Numbers indicate the number of codons inferred to be under diversifying or purifying selection by each method. Diversifying selection refers to codons with evidence of positive selection, whereas purifying selection refers to codons with evidence of negative selection.

| Gene | dN/dS | SLAC |  | MEME |  | FUBAR |  |
| --- | --- | --- | --- | --- | --- | --- | --- |
|  |  | Diversifying | Purifying | Diversifying | Purifying | Diversifying | Purifying |
| ORF1a | 0.379 | 0 | 61 | 5 | 0 | 16 | 162 |
| ORF1b | 0.179 | 0 | 55 | 1 | 0 | 2 | 192 |
| ORF3 | 0.215 | 0 | 0 | 0 | 0 | 1 | 6 |
| Spike (S) | 0.887 | 9 | 20 | 17 | 0 | 25 | 21 |
| Envelop (E) | 0.750 | 0 | 0 | 0 | 0 | 0 | 0 |
| Membrane (M) | 0.254 | 0 | 0 | 0 | 0 | 0 | 0 |
| Nucleocapsid (N) | 0.580 | 0 | 1 | 0 | 0 | 1 | 8 |

### Genomic signatures associated with PEDV population structure

We assessed the genomic regions that contributed most to the differentiation of the clusters inferred by DAPC by inspecting the sites with the highest loading values across the discriminant functions (Figure 6). In total, we identified 29 sites that strongly contributed to the structure of the clusters recovered by DAPC (Figure 7, Figure S2, S3). Each discriminant function revealed a distinct set of informative sites, indicating that different genomic regions contribute to different levels of separation among the groups. We then mapped each informative nucleotide site to its corresponding codon and translated these codons to identify the encoded amino acids. Because some of these sites were located within the same codon, the 29 sites corresponded to 26 distinct codons. 18 codons were located in the Spike protein, five in ORF1b, and three in ORF1a.

**Figure 7.**
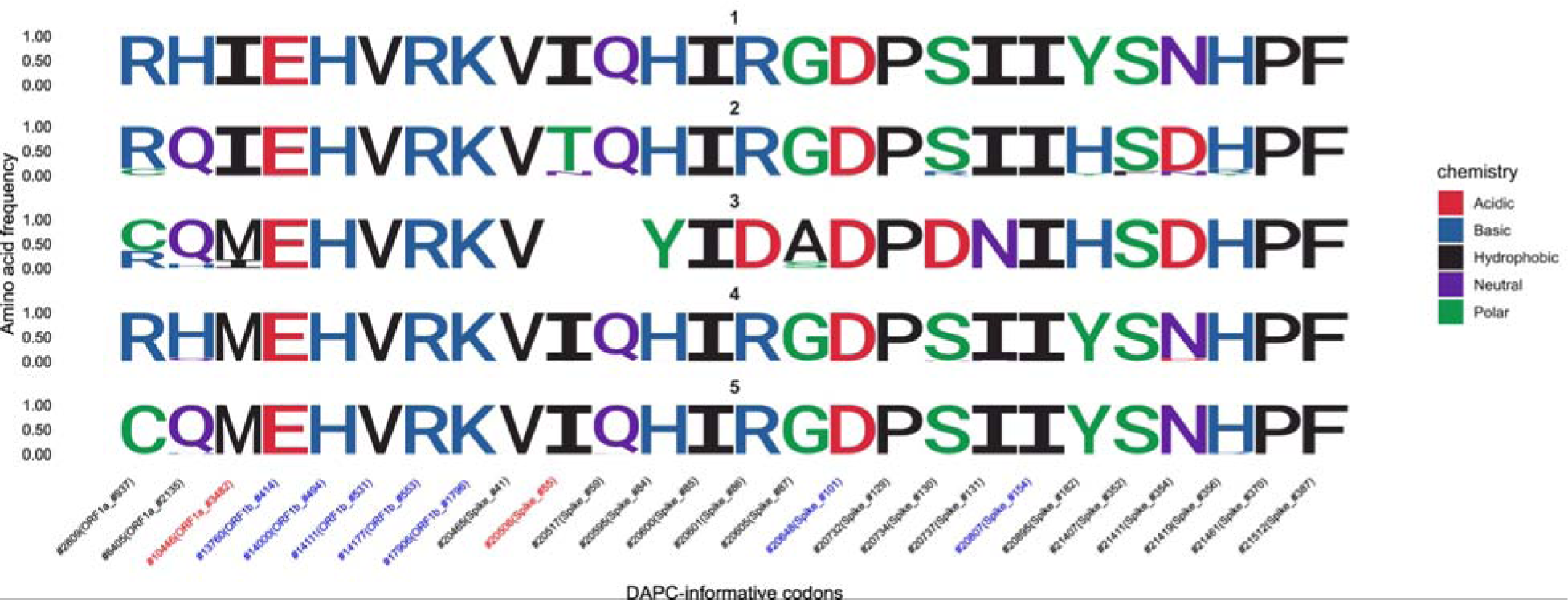
Amino acids encoded by codons containing DAPC-informative nucleotides sites associated with the genetic structure of the population. The sequence shows the frequency distribution of each amino acid encoded by codons containing nucleotide sites with the highest contribution to discrimination among groups inferred by DAPC. On the x-axis, numbers outside parentheses indicate nucleotide positions within the complete coding sequence, excluding UTRs; numbers inside parentheses indicate the corresponding amino acid positions within the indicated ORF. Sites indicated in blue correspond to those for which evidence of negative selection was detected by at least one method (Supplementary Table 1), whereas sites indicated in red correspond to those for which evidence of positive selection was detected by at least one method (Supplementary Table 1). The number above each sequence indicates the cluster to which it belongs. Amino acids are colored according to their biochemical class.

Next, we evaluated whether the DAPC-informative sites resulted in synonymous or nonsynonymous substitutions. Of the 26 informative codons, 13 involved synonymous substitutions and 13 involved nonsynonymous substitutions (Figure 7). Overall, nine of these codons also showed evidence of selection according to at least one of the methods previously applied. Seven codons were identified as being under purifying selection by at least one of the three selection inference methods (Supplementary Table 1), including five codons in ORF1b and two in the Spike gene. These results indicate that some of the sites contributing to population structure occur within functionally constrained regions, where genetic differences among clusters can accumulate despite the action of purifying selection.

Two DAPC-informative codons showed evidence of diversifying selection: codon 3482 in ORF1a (residue 199 in nsp6) and codon 55 in the Spike gene (Figure 7). Codon 3482 in ORF1a encoded either isoleucine or methionine. Clusters 1 and 2 predominantly encoded isoleucine at this position, whereas clusters 4 and 5 predominantly encoded methionine. Cluster 3 contained both amino acids, although methionine was the most frequent residue. Both amino acids are hydrophobic, representing a conservative substitution (Figure 7). The Codon 55 of the Spike protein was particularly relevant for cluster 2, which corresponds to the recent non-S-INDEL group. Most sequences in cluster 2 encoded threonine at this position, an uncharged polar amino acid, whereas the early non-S-INDEL clusters encoded isoleucine, a hydrophobic amino acid. In contrast, cluster 3, representing the S-INDEL strains, contained a deletion encompassing this region. Interestingly, the amino acid substitutions at Spike positions 182 and 353 were shared between clusters 2 and 3.

The previous analysis identified the substitutions associated with the genetic clusters inferred by DAPC. We next applied the same approach to the DAPC stratified by epidemiological period to determine whether the same genomic regions were also associated with the epidemiological periods (Figure 8, Figures S4 and S5). Only 11 informative sites were recovered in this analysis. Of these, 10 differed from those identified in the DAPC-inferred genetic clusters, with Spike codon 55 being the only site shared between the two analyses. Of the 11 informative codons, eight corresponded to synonymous substitutions and three to nonsynonymous substitutions. In addition, three codons, all located in ORF1a, showed evidence of purifying selection.

**Figure 8.**
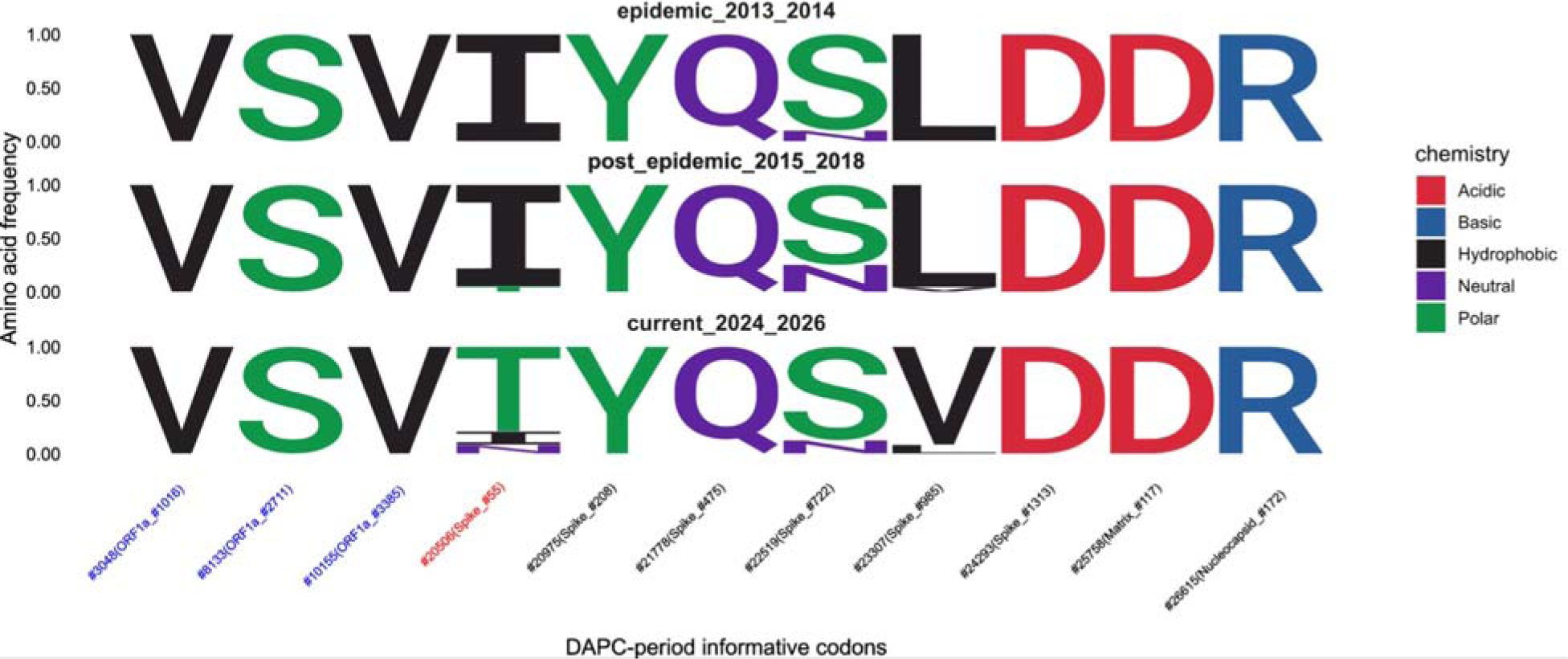
Amino acids encoded by codons containing DAPC-informative nucleotide sites associated with the genetic differentiation among epidemiological periods. The sequence logos show the frequency distribution of each amino acid encoded by codons containing nucleotide sites with the highest contribution to discrimination among epidemiological periods in the DAPC. On the x-axis, numbers outside parentheses indicate nucleotide positions within the complete coding sequence, excluding UTRs; numbers inside parentheses indicate the corresponding amino acid positions within the indicated ORF. Sites indicated in blue correspond to those for which evidence of negative selection was detected by at least one method (Supplementary Table 1), whereas sites indicated in red correspond to those for which evidence of positive selection was detected by at least one method (Supplementary Table 1). The label above each logo indicates the epidemiological period represented by that group. Amino acids are colored according to their biochemical class.

Taken together, these results indicate that the population subdivision inferred by DAPC is associated with distinct combinations of nucleotide substitutions rather than individual diagnostic mutations. Interestingly, Spike codon 55 emerged as the only amino acid position identified by both the genetic cluster-and epidemiological period-based analyses, distinguishing the contemporary non-S-INDEL strain from the remaining populations. In addition, a subset of the informative codons showed evidence of purifying selection, suggesting an association between selective constraint and the genomic differentiation identified by DAPC.

## Discussion

The introduction of PEDV into the United States in 2013 had a substantial impact on the North American swine industry and prompted considerable efforts to characterize viral genetic diversity, including the generation of a large number of complete genome sequences early after its introduction (7, 8, 13). After the epidemic period, sequencing efforts decreased, particularly for complete genomes, creating a substantial gap in our understanding of how the viral population continued to evolve after the early epidemic period (13). A recent study by Herrera da Silva et al. (13) partially addressed this gap using Spike (S1) sequences, showing that much of the diversity observed during the epidemic period was no longer detected in contemporary datasets and that only two clades appeared to persist in the current population, each with an apparently restricted geographic distribution. Specifically, Clade 1 appeared to be restricted to the Midwestern and Eastern regions whereas the Clade 2 was restricted to the Western United States (13). Although complete genomes were also analyzed in that study, those sequences had been collected only up to 2017, so the analysis of contemporary PEDV populations relied exclusively on S1 sequences. Therefore, the genetic diversity of contemporary PEDV populations at the whole-genome level remained largely unexplored. By generating complete PEDV genomes from samples collected between 2018 and 2026, our study adds important insights into PEDV diversification following the epidemic period. Our findings suggest that both recombination and natural selection have likely contributed to shaping the contemporary population structure of PEDV.

Despite breeding farm incidence remaining below 5% in recent years (13), the whole-genome analyses suggest that PEDV populations in the United States have undergone continuous diversification resulting in a pattern of genetic structuring over time. Our findings also suggest that the contemporary PEDV population is predominantly composed of a phylogenetically diverged group of viruses. However, viruses that are more genetically close to sequences from the early non-S-INDEL populations continue to circulate, apparently at low frequency. We observed a progressive increase in the genetic variability over time at the whole genome level, from the epidemic period to post-epidemic to current. This pattern was consistent across both the unsupervised and time-stratified DAPC analyses. A sampling gap for complete genomes remains for the 2019–2023 period. Consequently, it remains unclear whether this temporal structuring process occurred gradually or abruptly. Accessing archival samples collected during this window could provide important pieces for understanding the demographic dynamics of PEDV during this period.

The significantly negative Tajima’s D values observed during the epidemic and post-epidemic periods indicated an excess of low-frequency variants, a pattern consistent with rapid population expansion following a reduction in genetic diversity (54). This demographic signature is compatible with the known epidemiological history of PEDV in the United States, in which the introduction of a limited viral population likely resulted in a founder effect, followed by rapid epidemic expansion and the progressive accumulation of new genetic variation (10, 11, 13). In contrast, the contemporary PEDV sequences exhibited a negative but non-significant Tajima’s D value, under both analytical approaches (unsupervised DAPC and stratified by epidemiological period), indicating that the allele-frequency spectrum did not deviate significantly from neutral expectations. This result should be interpreted cautiously because substantially fewer complete genome sequences were available for the current period than for the preceding epidemiological periods, potentially reducing the statistical power to detect deviations from neutral expectations (54).

Our estimates of nucleotide diversity across the PEDV genome using a sliding-window approach revealed distinct hotspots of genetic variability within ORF1a, particularly in regions encompassing nsp2 and the papain-like protease (PL-Pro), and within Spike gene, predominantly in the S1 subunit. Although the function of nsp2 remains poorly characterized in PEDV and is not completely understood even in other coronaviruses (58), available evidence suggests that it may participate in interactions associated with the viral replication complex and in the modulation of the intracellular environment to support viral replication (59). Recent evidence identified nsp2 as a virulence determinant in PEDV (58). Although nsp2 is not essential for viral replication, its deletion reduces viral replication *in vitro* and produces an avirulent phenotype in neonatal piglets by reducing the virus’s ability to antagonize type I interferon responses through TBK1 degradation (58). Therefore, the high variability observed within nsp2 identifies this poorly characterized protein as an important candidate for future functional investigation.

The functions of PL-Pro have been more extensively characterized (59). PL-Pro is a multifunctional protein with an essential role in viral polyprotein processing and the release of mature nonstructural proteins, and in several coronaviruses it also exhibits deubiquitinating and deISGylating activities capable of modulating host innate immune responses, including interferon-associated pathways (59–61). Because of its essential role in processing the viral polyprotein and releasing the mature nonstructural proteins required for viral replication, PL-Pro has attracted considerable interest as a target for antiviral drug development in coronaviruses (60–63).

The Spike protein mediates interactions involved in viral attachment and entry and contains important targets of the host antibody response (64). Spike is also the principal antigen included in current PEDV vaccine strategies, and the elevated variability observed within S1 may have important implications for antigenic diversity and vaccine performance (64). Taken together, the elevated variability identified within nsp2, PL-Pro, and S1 illustrates how whole-genome sequencing not only provides a broader view of viral evolutionary processes but also represents a fundamental tool for understanding the genetic basis of viral phenotypes and identifying genomic regions that may contribute to vaccine development and updating, as well as to antiviral drug development (65–67).

Half of the informative sites supporting the differentiation among DAPC clusters involved synonymous substitutions, whereas the remaining half corresponded to nonsynonymous substitutions. Most of these informative sites also showed evidence of purifying selection, whereas only two were associated with diversifying selection, indicating that natural selection likely contributes to the temporal structuring of PEDV populations. Our selection analyses indicated that two sites contributing to the subdivision into the five clusters inferred by DAPC were under diversifying selection: site 3482 of ORF1a, corresponding to position 199 in nsp6, and site 55 of the Spike protein.

The nsp6 protein, together with nsp3 and nsp4, is involved in the formation of coronavirus replication organelles (68–70). The nsp6 is also involved in the modulation of autophagy. During PEDV infection, autophagy has been shown to exert a proviral effect by promoting viral replication (71). Previous studies demonstrated that pandemic PEDV is more efficient at modulating autophagy than classical strains, and that this is correlated with increased proliferation in intestinal porcine epithelial (IPEC-J2) cells (72). We identified an I-to-M substitution at nsp6 site 199 (site 3482 in ORF1a). This substitution was present in cluster 1 (part of the early non-S-INDEL strain) and cluster 2 (recent non-S-INDEL), whereas cluster 3 (S-INDEL) also presented this substitution, but methionine was the predominant amino acid. To the best of our knowledge, no information is available regarding the function of nsp6 residue 199. The substitution involves two amino acids of similar size and belonging to the same biochemical class, suggesting conservation of the biochemical properties at this site.

Site 55 in the Spike protein was informative for population subdivision in both the unsupervised DAPC clusters and the analysis based on epidemiological periods. The Spike protein is involved in host-cell attachment and entry (64). Site 55 is located within the N-terminal domain of the S1 subunit. One of the characteristic deletions found in North American S-INDEL strains spans residues 55–59, whereas these residues are retained in non-S-INDEL strains (8, 10). In previous studies using reverse genetics, partially overlapping deletions within this region were experimentally characterized. Deletions of residues 56–57 or 56–60 were associated with reduced pathogenicity (73). Among the sequences analyzed here that retained this region (i.e., non-S-INDEL strains), we identified variation at residue 55. The contemporary non-S-INDEL cluster was characterized by an I-to-T substitution involving amino acids with distinct physicochemical properties. Although the specific function of residue 55 remains unknown, its identification as a DAPC-informative site under diversifying selection, together with its location within a pathogenicity-associated region of the Spike N-terminal domain, highlights this residue as a candidate that should be prioritized for future functional characterization.

Finally, our time-scaled phylogenetic analyses reinforce previous evidence suggesting that only two contemporary Spike clades (Clade 1 and 2) are currently circulating in the United States. Interestingly, the evolutionary histories inferred from whole genomes and the Spike protein diverge. The contemporary whole genomes clustered predominantly within a single monophyletic clade divergent from the pre-2017 strains, with the exception of only two contemporary sequences that were more closely related to the early non-S-INDEL strains. The S1 phylogeny revealed that Clade 1, which was previously thought to be restricted to the Midwest and Eastern United States (13), was also detected circulating in the western United States. In contrast, Clade 2 was not detected outside the western United States. More extensive sampling may provide further evidence as to whether Clade 2 is in fact restricted to the western United States, while also expanding our understanding of viral dispersal among these regions.

PEDV elimination has been identified as a priority for the U.S. swine industry (9). Active genomic surveillance can help characterize viral population evolution and, when supported by representative sampling, estimate the regional prevalence and spatial distribution of circulating clades. Combined with epidemiological data, it can also help reconstruct dispersal routes between regions and identify potential source regions where containment and elimination efforts could be intensified. Such information has considerable potential to support regional exclusion strategies aimed at preventing the introduction or re-introduction of viral clades that are absent from a given region. Furthermore, integrating genetic data with field observations could help identify candidate genetic markers associated with phenotypes such as transmissibility and disease severity, supporting the identification of viral variants of concern following validation.

Nevertheless, some limitations remain. All sequences generated in this study were classified as non-S-INDEL. Samples were obtained exclusively from clinical cases, introducing a sampling bias, as S-INDEL strains tend to cause milder clinical signs and thus may have not been captured by our study (74). Indeed, S-INDEL viruses continue to circulate in the United States (75). Therefore, contemporary PEDV diversity in the United States may be greater than captured in this study. Our dataset was temporally unbalanced, with substantially greater sampling intensity during the epidemic period, and a considerable sampling gap remains between 2019 and 2023. Together, these limitations prevented us from drawing robust conclusions about the demographic processes affecting the contemporary PEDV population and from determining whether the observed temporal differentiation occurred gradually or through a more abrupt transition. In addition, geographic patterns were not explored in greater detail because of data sensitivity and the need to preserve the anonymity of participating farms.

This study represents a significant effort to characterize the evolutionary dynamics of PEDV from a whole-genome perspective, advancing our understanding after a nine-year period during which no new complete PEDV genomes had been generated. Overall, our findings demonstrate that PEDV has continued to diversify despite the substantial decline in disease incidence since the epidemic period. Our analyses indicate that most contemporary PEDV genomes cluster within a divergent clade and that the contemporary viral population is genetically more diverse than the populations that circulated during the epidemic and post-epidemic periods. In addition, the identification of genomic regions with elevated variability, including nsp2, PL-Pro, together with specific residues in nsp6 and Spike under diversifying selection, highlights candidate regions and sites for future studies investigating the mechanisms underlying viral adaptation, pathogenicity and virus–host interactions, with potential implications for the development of antiviral strategies. Ultimately, continued whole-genome surveillance will be essential for monitor ongoing viral evolution, inform regional exclusion strategies aimed at preventing the introduction of viral clades that are currently absent and supporting development of vaccines, antiviral strategies, thereby contributing to more effective PEDV control and eradication programs.

## Author Contributions

Conceptualization: J.P.H.S., and K.V.; methodology: J.P.H.S, and K.V.; Investigation: J.P.H.S., and K.V.; database curation: J.P.H.S, M.K., X.Y., A.R., F.V., J.M.,M.P.K., J.P., Q.Z., and C.A.C.; formal analysis: J.P.H.S., and K.V.; original draft writing: J.P.H.S.; review and editing: J.P.H.S., K.V., I.A.D.P., M.K., J.M., M.T., A.R., N.P.; Supervision: K.V.; Funding acquisition: K.V. and J.P.H.S., All authors have read and agreed to the published version of the manuscript.

## Data availability statement

The sequences used in this study are listed in the supplementary file sequence_list.xls. The complete-genome sequences generated in this study have been deposited in GenBank under accession numbers XXXXX–YYYYY. Portions of the dataset are privately owned by the production systems and are subject to access restrictions. These data may be made available upon reasonable request to the corresponding author, contingent upon permission from the respective production systems.

## Acknowledgments

The authors thank the U.S. swine production systems that voluntarily collected and provided samples for sequencing, as well as the participants of the Morrison Swine Health Monitoring Project (MSHMP) for their contributions to this study.

